# Gentamicin impairs nephron development in iPSC-derived kidney organoids and disrupts proximal tubular lysosomal function

**DOI:** 10.64898/2026.10.28.762495

**Authors:** E. Sendino Garví, W. Pioruńska, L. Böhl, R.W. Sparidans, R. Masereeuw, A.M. van Genderen

## Abstract

Gentamicin is an aminoglycoside antibiotic used to treat Gram-negative bacterial infections. However, its clinical use is limited by nephrotoxicity, affecting approximately 10 - 20% of treated patients. Nevertheless, gentamicin remains widely used due to its high potency, low cost, and low incidence of bacterial resistance and hypersensitivity. Importantly, gentamicin crosses the placental barrier and is administered to neonates, including preterm infants in whom nephrogenesis is still ongoing, raising concerns about its effects on kidney development. Current knowledge of this risk relies largely on rodent studies, which fail to fully recapitulate human kidney physiology. Here, we developed human-relevant *in vitro* models to investigate the impact of gentamicin on kidney development and to unravel the dynamic cellular responses underlying gentamicin-induced impairment.

Kidney organoids differentiated from induced pluripotent stem cells (iPSC) and magnetically sorted proximal tubule cells were used to model immature kidney and immature proximal cells, respectively. Organoids were continuously exposed to gentamicin (0.25 - 1.5 mM) throughout development (d7+18), while megalin-expressing proximal tubule cells were exposed for 24 hours to explore the cellular mechanisms underlying gentamicin toxicity. Intracellular concentrations of the gentamicin congeners C1, C1a, and C2/C2a/C2b were quantified by LC-MS/MS. Gentamicin exposure impaired organoid growth, and nephron formation and maturation. Additionally, gentamicin induced lipid accumulation and reduced lysosomal area in proximal tubule cells, indicating disrupted cellular metabolism and homeostasis. In conclusion, iPSC-derived kidney organoids provide a suitable platform to further investigate gentamicin’s impact on kidney development and studying co-treatments to mitigate its adverse effects.

## 1. INTRODUCTION

Gentamicin is a potent aminoglycoside antibiotic that has been in clinical use since 1970 and is primarily used to treat serious infections caused by Gram-negative bacteria, including endocarditis, sepsis, pneumonia, meningitis, urinary tract infections, and bone infections (Phillips et al. 1977). Additionally, when combined with other antibiotics such as β-lactamase, gentamicin is also effective against certain Gram-positive pathogens (Wells et al. 1992). Gentamicin gained widespread clinical use due to its high potency, relatively low cost, and low incidence of bacterial resistance and hypersensitivity (Hodiamont et al. 2022). However, its application is limited by well-documented toxicities, including nephrotoxicity, which appears in approximately 10-20% of patients receiving gentamicin treatment (Quiros et al. 2011). Early signs of gentamicin nephrotoxicity are detected through urinary biomarkers such as clusterin, kidney injury molecule-1 (Kim-1), cystatin C, and neutrophil gelatinase-associated lipocalin, pointing towards acute tubular injury (Klementa et al. 2025; Lei et al. 2018).

Although its use declined as safer antibiotics became available, gentamicin has regained clinical importance in the context of rising antibiotic resistance (Wargo and Edwards 2014). Nowadays, it is frequently administered in combination therapies or as a treatment of last resort when other options fail. Despite gentamicin is classified by the FDA as a pregnancy category D drug (Nguyen et al. 2025), which corresponds to drugs that have been showed to cause risk to the fetus but their potential benefits might outweigh the risks, it continues to be administered to pregnant women as a last-resort treatment when no safer alternatives are available. Gentamicin readily crosses the placental barrier, and studies have demonstrated that it accumulates in fetal kidneys (Gilbert et al. 1990). Additionally, in clinical practice, gentamicin is also widely used neonates and premature infants (Serane et al. 2009), in whom nephrogenesis is still ongoing, particularly to treat acute bacterial meningitis and neonatal sepsis (Chattopadhyay 2002; Fjalstad et al. 2013). Neonates are particularly vulnerable to gentamicin toxicity due to pharmacokinetics since, compared with adults, neonates exhibit reduced clearance and prolonged plasma half-life of gentamicin (Chattopadhyay 2002; Fjalstad et al. 2013; Lingvall et al. 2005; Lulic-Botica et al. 2014), leading to higher intracellular accumulation and prolonged exposure. Together, these clinical and developmental considerations underscore the urgent need to investigate how gentamicin impacts kidney development.

Although the exact mechanisms of gentamicin nephrotoxicity are not yet fully understood, available data, mostly derived from animal studies, provide insights into the possible pathways. Gentamicin is not a single molecule but a mixture of structurally related congeners, principally C1, C1a, C2, C2a and C2b, whose relative proportions vary between manufacturers and production batches. Gentamicin is a cationic drug that interacts with the negatively charged phosphoinositide components at the proximal tubule brush border membrane (8). It is internalized into proximal tubule cells primarily via receptor-mediated endocytosis involving the megalin-cubilin complex. Once taken by the cells, gentamicin accumulates in lysosomes, the Golgi apparatus, and the endoplasmic reticulum (Balakumar et al. 2010; Wargo and Edwards 2014). Within lysosomes, gentamicin disrupts phospholipid metabolism, resulting in phospholipidosis (Hostetler and Hall 1982; Wargo and Edwards 2014). Furthermore, when intra-lysosomal drug concentrations exceed a critical threshold, lysosomal membranes become destabilized, leading to permeabilization and release of gentamicin, as has been shown in animal studies (Randjelovic et al. 2017; Watanabe 1978), along with lysosomal proteases such as cathepsins, into the cytosol (Klementa et al. 2025).

In addition to its lysosomal effects, gentamicin released in cytosol directly targets mitochondria, impairing the respiratory chain, reducing ATP production, and generating reactive oxygen species (ROS), including superoxide and hydroxyl radicals. The resulting oxidative stress activates the intrinsic apoptosis pathway (Randjelovic et al. 2017). Collectively, these mechanisms drive proximal tubule cell injury and underline the nephrotoxic effects of gentamicin. While research has primarily focused on gentamicin’s toxic effects in mature kidneys, its impact during kidney development remains poorly understood.

The current models to study the effects of gentamicin during kidney development do not fully recapitulate structure and function of the human kidney. Besides, animal models showed to have an aminoglycoside nephrotoxicity threshold 13- to 60-fold higher than humans (Rodríguez Salgueiro and González Núñez 2016). Kidney organoids derived from induced pluripotent stem cells (iPSC) have emerged as powerful tools to study human-relevant nephrotoxicity, as they are three-dimensional, self-organizing multicellular structures that recapitulate key aspects of human kidney microarchitecture and function (Nakanoh et al. 2025). It is known that current iPSC-derived kidney organoids are immature and do not fully recapitulate an adult human kidney (Nakanoh et al. 2025) and thus, hold great potential to study drug effects on developing kidneys.

In this study, we aimed to establish a robust fetal-like organoid model to investigate how gentamicin affects kidney development. In addition, we aimed to create a responsive *in vitro* platform using isolated proximal tubule-like cells to monitor real-time intracellular processes upon gentamicin exposure to gain better insight on early-onset molecular mechanisms and better understand the mechanisms behind gentamicin nephrotoxicity.

## 2. RESULTS

### 2.1 Gentamicin is taken up by kidney organoids and induces cytotoxicity

To evaluate the potential of kidney organoids to study gentamicin nephrotoxicity during development, we first assessed the presence of key nephron segments, including podocytes, proximal tubules and distal segments. After differentiation and maturation (Figure 1A), brightfield and immunostainings revealed that the organoids developed structures representative of key nephron segments (Figure 1B-D). We detected the presence of glomeruli (∼15%), proximal tubules (∼37%), and distal segments (∼48%), (Figure 1D-E), being the proximal segments (podocytes and proximal tubules) >50% of the organoid composition, confirming the presence of nephron-like populations within the organoids. We further evaluated the gene expression of key markers of the proximal tubule to assess their maturity. All Ct values ranged from 24.3 to 34.8 confirming that at the transcriptome level the organoids express the genes associated with proximal tubule functioning and potentially highlighting a lower maturation state of the proximal tubules (Figure 1F).

**Figure 1.**
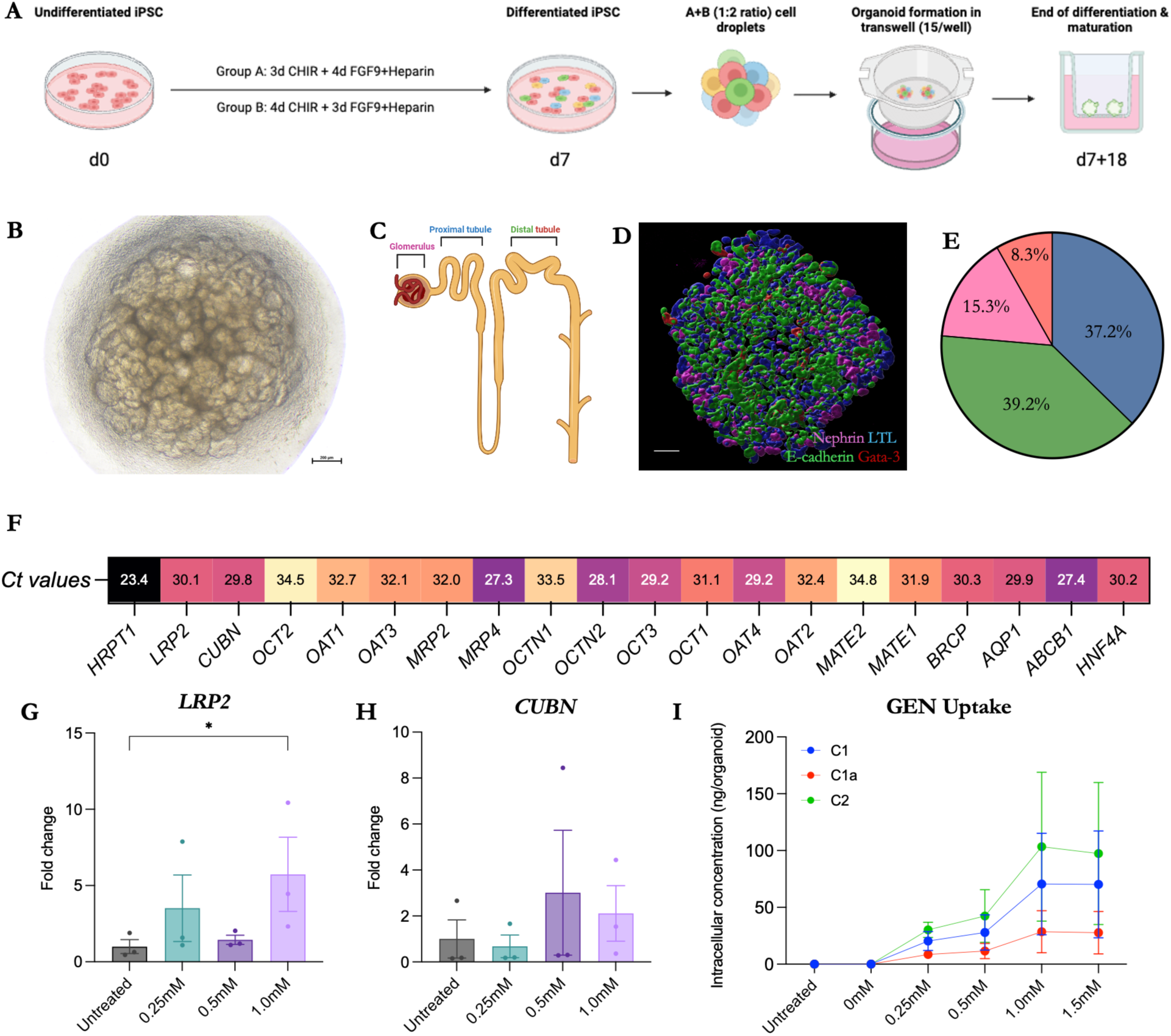
Kidney organoids internalize and are sensitive to gentamicin. **A)** Schematic representation of the differentiation and maturation process from iPSC to kidney organoids. **B)** Representative image of a kidney organoid at the end of the differentiation and maturation process at d7+18. Scale bar: 200 µm. **C)** Schematic representation of a nephron. **D)** Representative immunostaining rendered image from a kidney organoid at the end of the differentiation and maturation process at d7+18, in which nephrin appears depicted in pink as a marker of podocytes, Lotus tetragonolobus lectin (LTL) in blue as a marker for proximal tubules, and E-cadherin together with GATA-3 in green and red as markers for distal segments of the nephron, respectively. Scale bar: 200 µm. **E)** Diagram depicting the volume (as a percentage relative to all segments added together) of each nephron marker. **F)** Heatmap depicting the average Ct values (measured by qPCR) of several markers for proximal tubules in the untreated kidney organoids. **G & H)** Graphical presentation of the gene expression (measured by qPCR) of *LRP2* and *CUBN*, depicted as the fold-change of the gentamicin-treated organoids to the untreated control. **I)** Graphical presentation of the intracellular concentration of the three measured gentamicin congeners (C1, C1a, and C2/C2a/C2b) with increasing concentrations of gentamicin treatment. (N=3, in which each biological replicate is a kidney organoid of a different differentiation process, and each technical replicate is a different organoid within the same differentiation process for each condition. *= p<.05, **= p<.01).

Megalin (encoded by *LRP2*) and cubilin (encoded by *CUBN*) work together as a complex that is necessary for gentamicin uptake into the proximal tubule cells. Our transcriptomic analysis showed that our organoids express both *LRP2* and *CUBN* (Figure 1G-H) and that *LRP2* expression increased by 6-fold after exposure to 1 mM gentamicin (Figure 1G). In parallel, we measured the intracellular levels of gentamicin congeners (C1, C1a, and the sum of C2/C2a/C2b) following exposure to increasing concentrations of the drug to evaluate their uptake capabilities. We observed a concentration-dependent trend of gentamicin uptake, confirming that the organoids actively accumulate the drug, particularly C2/C2a/C2b (Fig 1I). Overall, these results confirm the kidney-like identity of the organoids and their ability to internalize and potentially respond to gentamicin treatment.

### 2.1 Continuous gentamicin exposure leads to impaired growth and low nephron count

To evaluate whether gentamicin exposure during development influenced the kidney organoids growth, the organoid size was measured at each day of the treatment starting three days after organoid seeding (Figure 2A). Over time, treatment with gentamicin prevented normal growth in a dose-dependent manner (Figure 2 B-C). While organoids treated at lower concentrations (0.25-0.5 mM) showed similar growth when compared to untreated controls, continuous treatment with 1 mM or 1.5 mM resulted in growth impairment (Figure 2B-C). To further evaluate whether the growth impairment was due to an overall organoid effect or a nephron segment-specific reduction, we measured the volumes of the different nephron segments (Figure 2D). At lower concentrations (0.25 mM), a somewhat increased presence of nephron segments (Figure 2D-E) was observed whereas exposure to higher doses led to a progressive reduction in nephron segments with the proximal tubules being the most affected and an almost complete loss of structures at 1.5 mM compared to the untreated control (Figure 2D-E). Structurally, we observed a loss in nephron segments continuity in gentamicin-treated organoids (0.5-1.5 mM), as evidenced by the presence of more unstained regions and isolated structures between the different segments, when compared to the untreated control.

**Figure 2.**
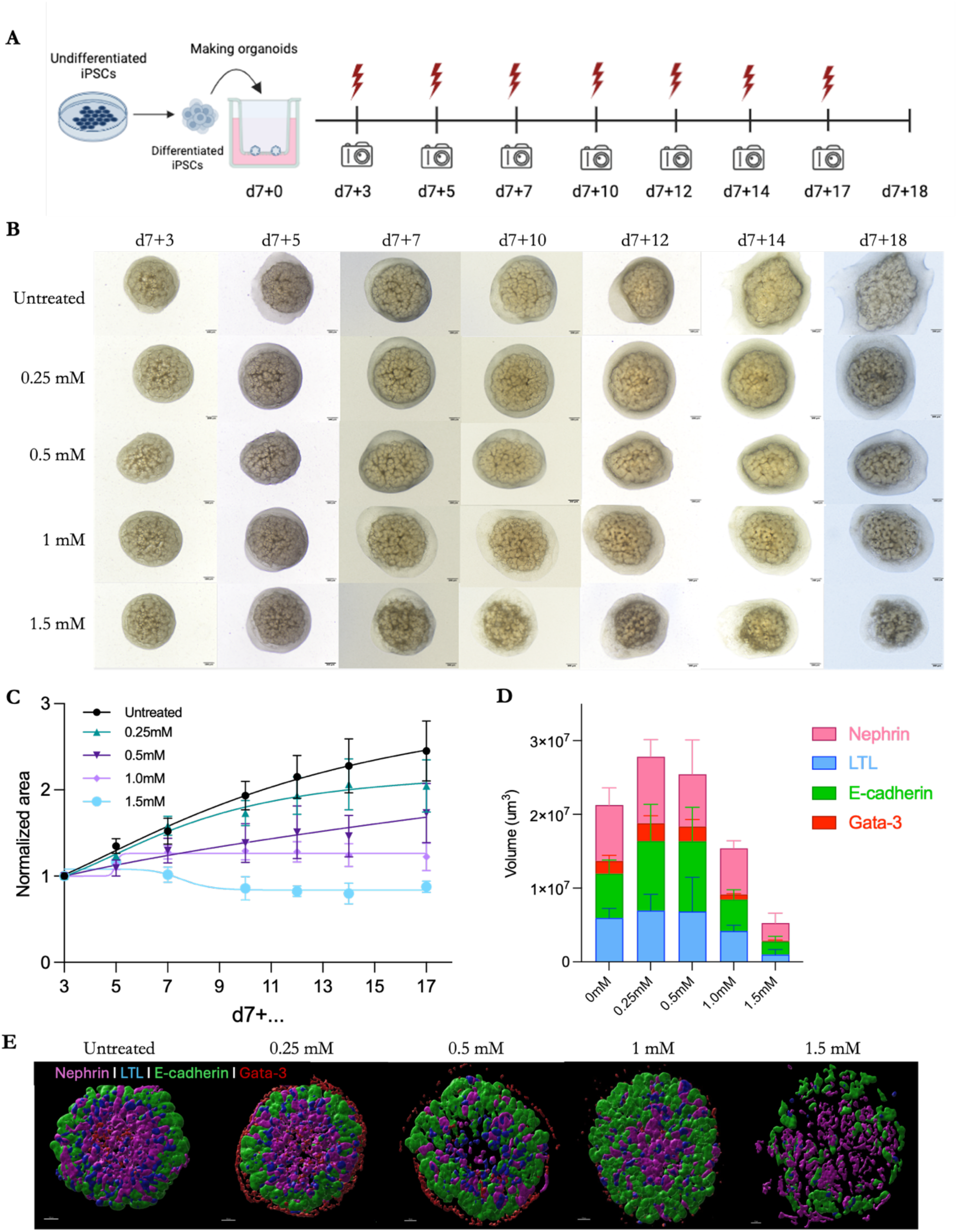
Continuous gentamicin exposure leads to impaired growth and low nephron number. **A)** Schematic representation of the gentamicin treatment timepoints during the differentiation and maturation process of the kidney organoids. **B)** Representative brightfield images of kidney organoids from the first day of gentamicin treatment (d7+3) up to the end of the differentiation and maturation process (d7+18). Brightfield images were taken every 2-3 days corresponding to the treatment days. **C)** Graphical representation of the growth of the kidney organoids measured as the area from brightfield images over the course of the treatment. **D)** Graphical presentation of the volume (in mm^3^) of the different nephron segments within the kidney organoids treated continuously with increasing doses of gentamicin. **E)** Representative immunostaining rendered images from kidney organoids exposed to increasing concentrations of gentamicin during the differentiation and maturation process. Scale bar: 200 µm. (N=3, in which each biological replicate is a kidney organoid of a different differentiation process, and each technical replicate is a different organoid within the same differentiation process for each condition.

Given that gentamicin preferentially accumulates in proximal tubules (19), this nephron segment is expected to be the most affected after gentamicin exposures. We analyzed the sphericity of the proximal tubules under the assumption that more spherical morphologies correspond to either to less mature or more structurally impaired tubules (Figure 3). Qualitatively, we observed impaired proximal tubule structures starting at 0.5 mM (Figure 3A), but our results only showed increased sphericity at 1.5 mM, when compared to the other treatment conditions (Figure 3B). Furthermore, gene expression analysis of kidney developmental markers (*WT1, SIX2,* and *HNF4a*) showed a trend towards concentration-dependent upregulation, with *WT1* expression at 1 mM being the strongest effect when compared to the untreated control (Figure 3C-E). Overall, these results suggest that gentamicin treatment affects organoid growth and that this reduction in growth at higher concentrations is driven by the loss of nephron segments, particularly the reduction in presence and maturation of proximal tubules.

**Figure 3.**
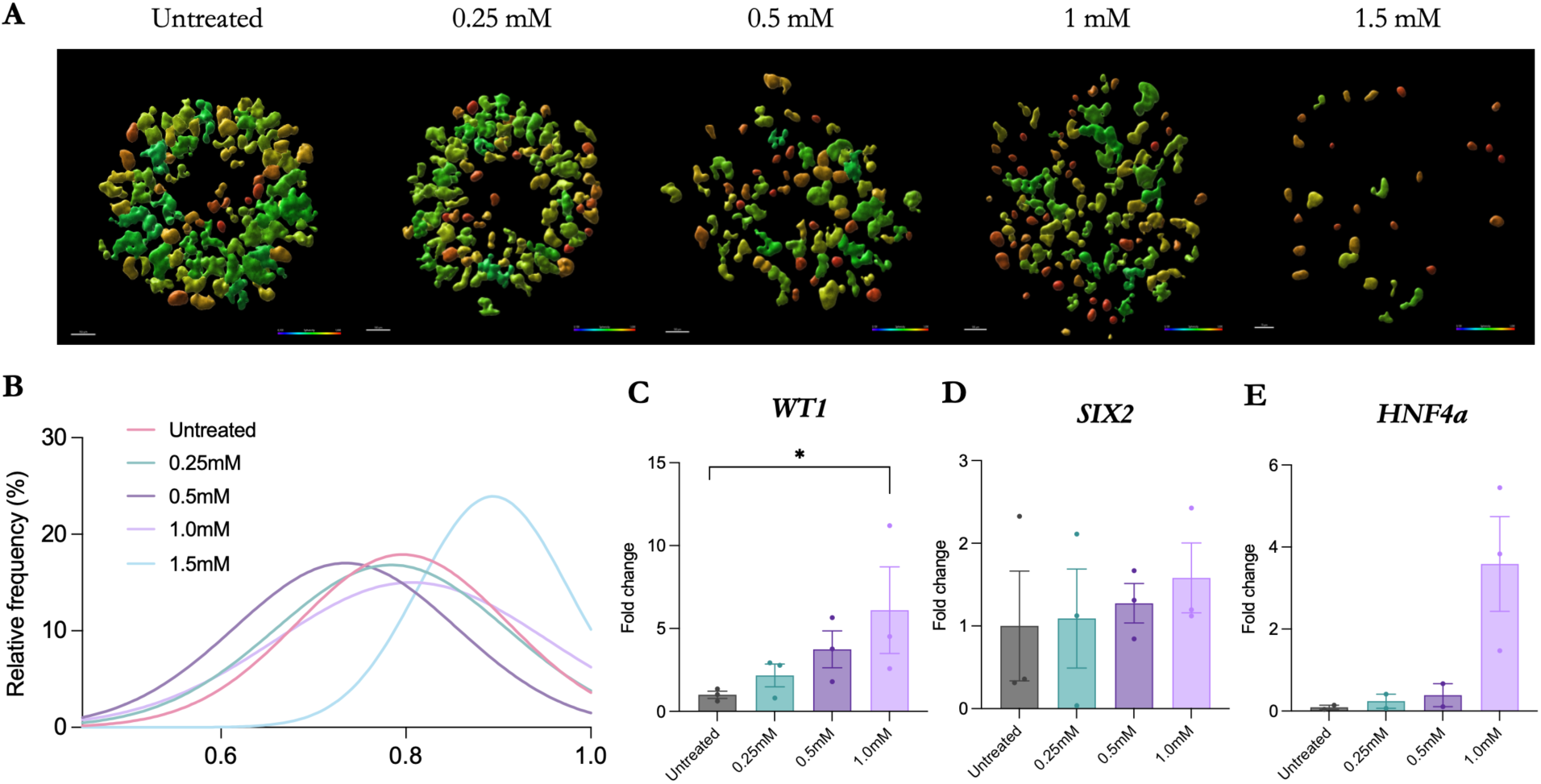
Continuous gentamicin exposure leads to organoid immaturity and proximal tubule loss. **A)** Representative immunostaining rendered images from the proximal tubules of the kidney organoids exposed to increasing doses of gentamicin during the differentiation and maturation process. Scale bar: 200 µm. **B)** Graphical representation of the sphericity of the proximal tubule structures within the kidney organoids. **C-E)** Graphical representation of the gene expression (measured by qPCR) of *WT1*, *SIX2*, and *HNF4a* depicted as the fold-change of the treatment groups to the untreated control, in kidney organoids treated with increasing concentrations of gentamicin during the differentiation and maturation process. (N=3, in which each biological replicate is a kidney organoid of a different differentiation process, and each technical replicate is a different organoid within the same differentiation process for each condition. *= p<.05).

### 2.1 Kidney organoids show the presence of early injury markers upon Gentamicin exposure

To investigate whether gentamicin treatment during kidney development induces early-onset and/or late-onset kidney injury, we evaluated the gene expression of early injury markers (*KIM-1, NGAL,* and *CST3*), as well as the presence of fibrosis (Figure 4). To evaluate whether gentamicin-induced injury is segment-specific or a generalized effect, we used *KIM-1* as a marker for proximal tubule injury, *NGAL* as a marker for distal segments injury, and *CST3* as a marker for overall organoid stress. Our transcriptomic analysis showed no significant changes for *KIM-1* (Figure 4A), suggesting proximal tubule-specific injury is less apparent or insufficient, or point towards less mature proximal tubules. A trend towards upregulation of *NGAL* was detected, particularly at 1 mM when compared to the untreated control (Figure 4B), suggesting that the distal segments might reach their stress threshold at higher doses. Lastly, we observed a consistent concentration-dependent upregulation trend in *CTS3*, reaching ∼2-fold increase when compared to control (Figure 4C). Overall, these results are suggestive of a global whole-organoid stress response rather than a segment-specific injury.

**Figure 4.**
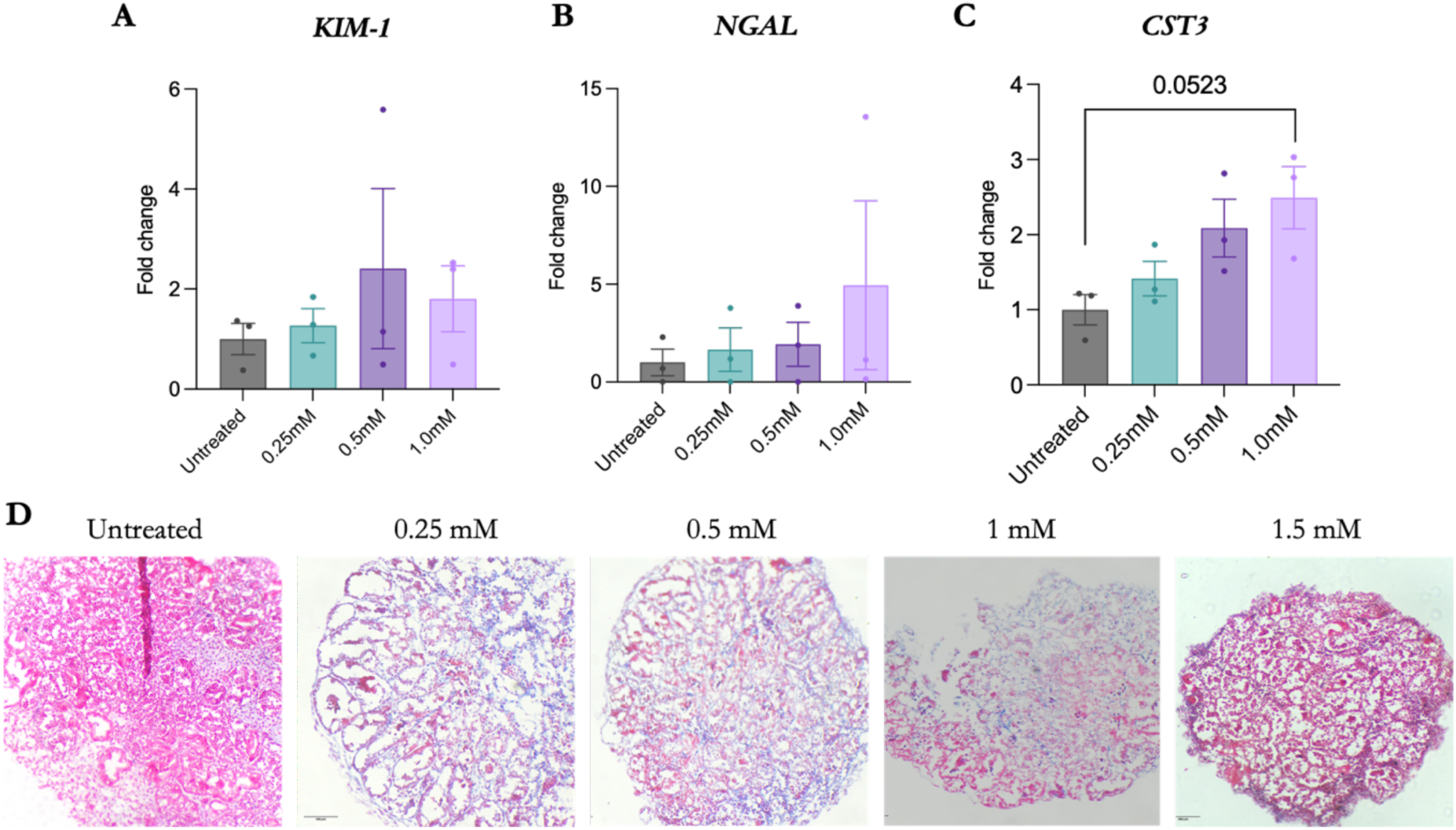
Kidney organoids show the presence of early injury markers and tubular necrosis upon Gentamicin exposure. **A-C)** Graphical representation of the gene expression (measured by qPCR) of *KIM-1*, *NGAL*, and *CST3* depicted as the fold-change of the treatment groups to the untreated control. **D)** Representative images of kidney organoids treated with increasing gentamicin doses during the differentiation and maturation process stained with Azan Trichrome. Structures in pink and blue depict non-collagenous and rich in collagen areas, respectively. Scale bar: 200 µm. (N=3, in which each biological replicate is a kidney organoid of a different differentiation process, and each technical replicate is a different organoid within the same differentiation process for each condition.

To evaluate whether gentamicin exposure during kidney development leads to fibrosis, we quantified the presence of collagenous tissue across different concentrations (Figure 4D). Despite we did not detect a significant difference amongst the different gentamicin exposures, we observed progressive necrosis in tubular structures across the organoids, starting at 0.25 mM, leading to almost complete loss of discernible tubules at higher concentrations.

### 2.1 Gentamicin prevents lysosomal formation and promotes lipid accumulation in proximal tubule like cells

To elucidate the acute effects of gentamicin exposure on the proximal tubule, we isolated proximal tubule cells (using magnetic beads towards LTL) (Figure 5A) from the kidney organoids at the end of the differentiation protocol and expose them to gentamicin for 24h. We first confirmed that isolated LTL+ cells retained their proximal tubule identity, as proven by LTL and megalin protein expressions (Figure 5B). We further confirmed that sorted cells were functionally capable of internalizing the sum of congeners C2/C2a/C2bdose-dependently, with particularly accumulation at high concentrations (1-1.5 mM) (Figure 5C). Over increasing gentamicin concentrations, the expression of megalin decreased over time while LTL+ signal remained stable (Figure 5D-E), suggesting that the loss of megalin is due to the treatment and not because a loss of proximal tubule identity of the cells.

**Figure 5.**
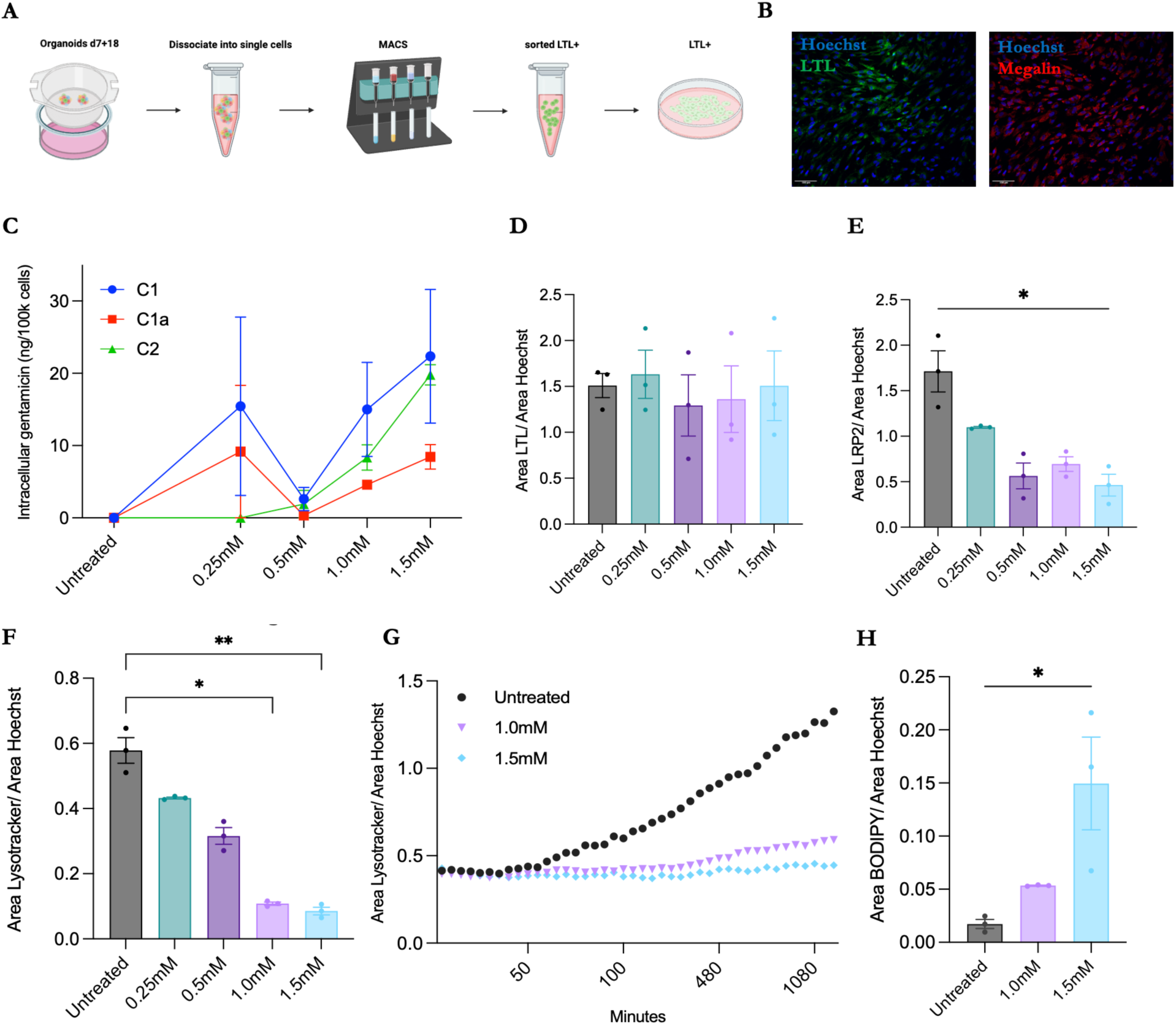
Gentamicin might prevent lysosomal formation and promote lipid accumulation in proximal tubule like cells. **A)** Schematic representation of the MACS sorting of LTL+ cells from the kidney organoids at d7+18. **B)** Representative immunostaining images of sorted proximal tubule like cells at passage 3 in which LTL and megalin are depicted in green and red, respectively. Scale bar: 100 µm. **C)** Graphical presentation of the intracellular doses of the three main gentamicin congeners (C1, C1a, and C2/C2a/C2b) with increasing concentrations of gentamicin treatment in proximal tubule like cells. **D-F)** Graphical presentation of protein expression (measured by area of LTL, megalin, and Lysotracker, respectively, over the Hoechst area). **G)** Graphical presentation of the area of lysosomes (measured by Lysotracker) over a 24h period in proximal tubule like cells treated with gentamicin. **H)** Graphical presentation of the presence of lipid droplets (measured by area of BODIPY over the Hoechst area) in proximal tubule like cells after 24h of gentamicin treatment. (N=3, in which each biological replicate are LTL+ cells sorted from kidney organoids of a different differentiation process, and each technical replicate is a different well within the same experiment. *= p<.05, **= p<.01)

One of the earliest detectable structural abnormalities in gentamicin-induced kidney injury is the “foamy” appearance of the proximal tubules because of lysosomal phospholipidosis, an acquired lysosomal storage disorder where excessive amounts of phospholipids build up inside the lysosomes of proximal tubule cells. To investigate whether this phenomenon occurs in our model, we measured the area of the lysosomes and the presence of lipids in proximal tubule like cells treated with increasing doses of gentamicin (Figure 5F-H). After 24h of exposure, our results showed a significant reduction in lysosomes when cells were treated with 1 or 1.5 mM compared to the untreated control. To determine whether this reduction in lysosomes was due to the inhibition of de-novo lysosomal formation or the destruction of existing lysosomes, we measured the lysosomal area over 24h of 1-1.5 mM of gentamicin exposure (Figure 5G). At the start of the experiment and for the first 50 minutes, the lysosomal area in the treatment groups was comparable to the untreated control. Over time, the lysosomal area and variability steadily increased in the untreated control proximal tubule like cells, while it remained unchanged and less variable in the 1-1.5 mM treatment groups, suggesting that gentamicin might disrupt the formation of new lysosomes rather than destructing the pre-existing ones. Inside the lysosomes, gentamicin inhibits phospholipases A and C (Hostetler and Hall 1982), resulting in lipid accumulation. To investigate this, we quantified the presence of lipid droplets at 1-1.5 mM doses of gentamicin after 24h of exposure in the proximal tubule like cells (Figure 5H). We observed a significant increase in lipid droplet accumulation in both treatment conditions when compared to the untreated control, consistent with lysosomal-metabolic stress. Overall, these results suggest that gentamicin may disrupt lysosomal homeostasis in proximal tubule cells by preventing lysosome formation and promoting lipid accumulation, recapitulating early hallmarks of aminoglycoside-induced nephrotoxicity.

## 3. DISCUSSION

This study demonstrates that iPSC-derived kidney organoids are a powerful model to investigate gentamicin-induced nephrotoxicity, revealing growth impairment and loss of nephron segments, particularly the proximal tubule, upon continuous exposure to the aminoglycoside. Mechanistically, gentamicin nephrotoxicity likely progresses from proximal tubule uptake via impaired lysosomal metabolism leading to lipid accumulation. Clinically, intravenous gentamicin dosing typically ranges from 5 to 7 mg/kg body weight (Choi et al. 1999), corresponding to serum concentrations of0.84-1.17mM, in adults, consistent with the concentrations used in this study (0.25 to 1.5 mM). In neonates (32-36 ± 6 weeks gestational age), a 4 mg/kg dose frequently resulted in serum levels outside the therapeutic range (Allegaert 2009; Serane et al. 2009), indicating altered pharmacokinetics relative to adults and potentially increasing the susceptibility of the developing kidneys to gentamicin toxicity. However, the effects of gentamicin on the developing kidneys of neonates have not been studied, underscoring the value of our model as platform for assessing nephrotoxicity in the immature kidney. Our results demonstrate that organoids take up all gentamicin congeners (C1, C1a, and C2/C2a/C2b), indicating pharmacologically responsiveness and active drug accumulation. The sum of congeners C2/C2a/C2b reached the highest intracellular levels, consistent with previous *in vitro* and *in vivo* animal studies identifying C2 as the most nephrotoxic congener and preferentially accumulated in proximal tubule cells (Bulman et al. 2020).

Gentamicin exposure impaired kidney organoid development, reducing overall organoid size and nephron volume, with all four nephron segments affected, particularly the proximal tubule. Given that gentamicin preferentially accumulates in proximal tubule cells via megalin-cubilin-mediated endocytosis, a pathway restricted to this segment (Dagil et al. 2013), the broader structural impairments observed suggest that developing nephrons are sensitive to gentamicin beyond the proximal tubule. Gentamicin can also enter cells through endocytosis-independent transport, including TRP cation channels such as TRPV4 (Berrout et al. 2012; Bulman et al. 2020), which is predominantly expressed in more distal segments of the nephron. Consistent with TRPV4-mediated uptake, the distal segment marker, -3, was absent at 1.5 mM gentamicin. We also observed upregulation of key kidney developmental markers (*WT1*, *SIX2*, and *HNF4A*) following gentamicin treatment. *WT1*, essential for nephrogenesis and later restricted to mature podocytes (Guo et al. 2002; Wang et al. 2024), was significantly increased at 1.0 mM, suggesting impaired or delayed organoid maturation. *SIX2*, a nephron progenitor marker (Kobayashi et al. 2008), showed a similar dose-dependent increase, further supporting retention of an immature state. *HNF4A*, a regulator of proximal tubule differentiation (Yoshimura et al. 2023), followed the same dose-response pattern, potentially reflecting a compensatory response to proximal tubule injury. Consistent with this, proximal tubules displayed a more spherical morphology at the highest dose, indicating disrupted structural maturation.

To further determine whether gentamicin triggers segment-specific or global injury responses, we assessed *KIM-1*, *NGAL*, and *CST3* as markers of proximal tubule damage (Klementa et al. 2025), distal segment stress (Lei et al. 2018), and overall organoid injury (Lei et al. 2018), respectively. *KIM-1* expression remained unchanged across conditions, suggesting that proximal tubule injury was either insufficient to elicit a detectable transcriptional response or that immature proximal tubules in these organoids are less responsive to injury. *NGAL* expression increased only at 1 mM, suggesting that distal nephron segments may reach a stress threshold only under more severe conditions. *CST3*, showed the clearest pattern, with a robust and dose-dependent upregulation up to 1 mM, reaching approximately a two-fold increase relative to controls. Together, these marker profiles point toward a generalized, whole-organoid stress response rather than a strictly segment-restricted injury pattern.

Given the prominent gentamicin effects on proximal tubules observed in our organoids, and the classical attribution of the aminoglycoside nephrotoxicity to this nephron segment (Nagai and Takano 2014; Quiros et al. 2011), we isolated LTL+ cells to examine intracellular processes at higher resolution. The sorted LTL+ population maintained a stable proximal tubule identity, showing strong LTL expression and detectable megalin, a receptor often poorly expressed in commonly used proximal tubule cell lines such as HK-2 (Nagai and Takano 2014). Megalin levels decreased with increasing gentamicin concentration, consistent with the known mechanism whereby gentamicin binding triggers internalization of the megalin-drug complex, reducing receptor availability at the cell surface (Beenken et al. 2023; Keller et al. 2024). This receptor depletion may, in turn, drive compensatory transcriptional upregulation to restore transport capacity, a response observed in the kidney organoids as well. Furthermore, we observed a dose-dependent attenuation in lysosomal area expansion of over 24 h in gentamicin-treated cells. At high intracellular concentrations, gentamicin is known to induce lysosomal membrane permeabilization or rupture (Denamur et al. 2011; Mingeot-Leclercq and Tulkens 1999), releasing lysosomal contents, including the drug itself, into the cytosol. Such damage alters lysosomal pH, rendering lysosomes undetectable by pH-sensitive probes and producing the apparent reduction in lysosomal area observed here. An alternative, non-mutually exclusive explanation involves altered lysosomal dynamics: under normal conditions, cells continuously generate new lysosomes, but following gentamicin treatment, nascent lysosomes may instead fuse with drug-containing endosomes to form endo-lysosomes (Keller et al. 2024). This fusion would reduce the pool of individually detectable lysosomes, consistent with the known uptake route of gentamicin, which is enters via adsorptive endocytosis and accumulates as endocytic vesicles fuse with lysosomes (Balakumar et al. 2010). Altogether, our findings demonstrate that kidney organoids provide a versatile, human-relevant model for studying gentamicin-induced nephrotoxicity. The combination of whole-organoid analysis and isolated proximal tubule like cells offers mechanistic insight into both developmental impairments and lysosomal dysfunction, providing a foundation for future investigation of nephrotoxicity mechanisms and protective interventions.

## 4. CONCLUSIONS

In conclusion, our findings establish a novel, human-relevant platform for studying the effects of gentamicin on kidney development and proximal tubule function. We show that gentamicin impairs organoid growth, nephron maturation, and proximal tubule structure, while inducing global stress responses and lysosomal-metabolic alterations indicative of early aminoglycoside nephrotoxicity. Importantly, LTL+ cells isolated from organoids retain proximal tubule identity and robust megalin expression, making them a responsive model for studying drug uptake, lysosomal dysfunction, and intracellular stress pathways, features often poorly represented in conventional 2D proximal tubule models. Together, these results establish kidney organoids and their derived proximal tubule cells as a versatile system for assessing gentamicin-induced toxicity and investigating protective or therapeutic interventions in a human context.

## 5. MATERIALS AND METHODS

### 5.1 Reagents and antibodies

All media and their corresponding supplements used for cell culture and kidney organoid differentiation were obtained from Gibco, Life Technologies (Paisley, UK) unless stated otherwise. Suspension plates used for iPSC culture were coated with Geltrex® LDEV-Free hESC-Qualified Reduced Growth Factor Basement Membrane Matrix (#A1413302, Gibco Life Technologies, Paisley, UK) prior to passaging.

The primary antibodies used in this study were human anti-nephrin (#AF4269-SP, R&D Systems, Abingdon, United Kingdom) diluted 1:300, biotinylated lotus tetragonolobus lectin LTL (#B-1325, Vector Laboratories, Brunschwig Chemie, Amsterdam, The Netherlands) diluted 1:300, mouse anti-E—cadherin (#610181, BD Biosciences, Mississauga, ON, Canada) diluted 1:300, rabbit anti-GATA-3 (#5852S, Cell Signaling Technology, Leiden, The Netherlands) diluted 1:300, TIM-1 polyclonal antibody (#PA5-20244, Invitrogen, Waltham, MA, USA) diluted 1:25, rabbit anti-LRP2 (#ab76969, Abcam, Cambridge, UK) diluted 1:100. The probe used for monolayer assessment was Phalloidin 647 (#ab176759, Abcam, Cambridge, UK) diluted 1:1000. For detection of lipid droplets BODIPY 493/503 (#D3922, ThermoFisher Scientific, Waltham, MA, USA) probe diluted 1:1000 was used. All secondary antibodies were added at a final dilution of 1:400 and all were obtained from ThermoFisher Scientific (Breda, The Netherlands). The secondary antibodies used for detection were Alexa Fluor 647 donkey anti-sheep (#A-21448), streptavidin Alexa fluor 405 conjugate (#S32351), Alexa fluor 488 donkey anti-mouse (#A21202), Alexa Fluor 568 donkey anti-rabbit (#A10042), donkey anti-rabbit 647 (#A31573).

### 5.2 Cell culture - Kidney Organoids and LTL+

#### 5.2.1 iPSC Culture

iPSCs generated from erythroblasts were obtained from Radboud UMC via a material transfer agreement with The Stem Cell Technology and differentiation Center (SCTC). Cells were cultured in colonies on 6-well plates (Cat. No. 665 102, Cellstar, Greiner, 6-well suspension culture plates) (2 mL/well) using Essential 8 medium supplemented with E8 supplement and maintained at 37 °C in a humidified incubator with 5% CO₂. Complete culture medium (referred to as E8+) was changed daily. For weekend maintenance, medium volume was adjusted to 6ml/well. Prior to passaging, 6-well plates were coated with 2% Geltrex® LDEV-Free hESC-Qualified Reduced Growth Factor Basement Membrane Matrix in DMEM/F12 (1:1) and incubated for 1 hour at RT to allow gel formation. Colonies were washed twice with sterile PBS (Dulbecco’s Phosphate Buffered Saline, D8537, Sigma Aldrich) and dissociated using 1 mL of UltraPure 0.5 mM EDTA (#15575-038, Invitrogen, NY, USA) for 4 minutes at RT. After removing EDTA, E8+ medium was gently added to dislodge the colonies. Cells were passaged at a ratio between 1:3 and 1:12, depending on confluency and growth rate.

#### 5.2.2 Kidney Organoid Differentiation

Kidney organoids were generated from the protocol described by Jansen et al (20) with minor adaptations. iPSCs were first washed with sterile PBS, followed by enzymatic dissociation into single cells using TrypLE Select (1X) (#12563-029, Gibco, Life Technologies, Grand Island, NY, USA). Enzymatic activity was quenched by adding E8+ medium. The cell suspension was collected, centrifuged at 400 × *g* for 3 minutes, and resuspended in an appropriate volume of E8+ with (1:100) Revitacell medium. Cells were seeded and incubated at 37 °C in 5% CO₂ for 24 hours (d-1). After 24 hours, the medium was replaced with pre-warmed Essential 6 (E6) Medium supplemented with 6 μM CHIR-9902 (#4423, Tocris Bioscience, Bristol, UK) (d0). After 3 days of CHIR treatment, cultures were split into two groups (Group A and Group B) at a 1:2 ratio. Group A was cultured in E6 medium supplemented with Human Recombinant FGF-9 1:500 (#78161, Stem Cell Technologies, Vancouver, British Columbia, Canada) and heparin 1:1000 (#07980, Stem Cell Technologies, Vancouver, British Columbia, Canada) for 2 days (d3 and d4). Group B continued treatment with E6 + 6 μM CHIR-9902 for an additional 2 days (d3 and d4). After this period, both groups were further cultured in E6 medium containing Human Recombinant FGF9 (1:500) and heparin (1:1000) for an additional 2 days. iPSCs differentiated in 2D formed a confluent monolayer and were harvested by first washing with PBS, followed by incubating with TrypLE Select for 2-3 minutes at 37 °C to dissociate the monolayer into single cells. Enzymatic activity was neutralized by adding DMEM/F12 supplemented with 10% fetal calf serum (FCS) at twice the volume of TrypLE used. Cells from group A and group B were mixed at a 1:2 ratio (A:B). The mixed cell suspension was centrifuged at 400 rcf for 3 minutes and resuspended in E6 medium at a concentration of 100,000 cells/μL. Droplets of 0.2 μL of the cell suspension containing 20,000 cells each were dispensed into Transwell inserts, PET membrane (CLS3450-24EA, Corning, NY, USA), approximately 15 organoids were seeded per insert. E6 medium supplemented with 5 μM CHIR-99021 was added to the basolateral chamber, and cells were incubated at 37 °C for 1 hour to allow aggregation (d7+0). Following incubation, the medium was replaced with 1.2 mL of E6 supplemented with FGF9 (1:500) and heparin (1:1000). The kidney organoids were cultured for a total of 18 days, with medium changes performed every 2-3 days. For the first medium change (d3), E6 with FGF9 and heparin was used. From the second medium change onward, organoids were maintained in E6 supplemented with Human BMP-7 Recombinant Protein 1:2000 (#120-03P-100UG, Peprotech, Cranbury, NJ, USA) and epidermal growth factor (EGF) 1:1000 (#E9644-.5MG, Sigma Aldrich, St. Louis, MO, USA).

#### 5.2.3 Isolation of proximal tubule cells (LTL+)

At the end of the differentiation protocol (d7+18) kidney organoids were collected from the Transwells (1mL of PBS/Transwell insert) and were then centrifuged at 400 rcf for 1 minute. After removing the PBS, 300 µL of 0.25% trypsin were added to dissociate them into single cells. The cells were incubated with trypsin until a homogeneous viscous suspension was formed, mixing every 5 minutes. Next, separation buffer with DNAse (1:1) was added to degrade any DNA, and the mixture was incubated for 5 minutes at RT, with mixing every minute. This step was followed by a centrifugation at 500 rcf for 4 minutes. The viscous material was discarded, and the two steps (DNase treatment and centrifugation) were repeated. Cells were incubated for 15 minutes with Biotinylated LTL antibody at 4°C. Following incubation, 8 mL of separation buffer was added, and the suspension was centrifuged at 300 rcf for 4 minutes. This wash step was repeated twice. After discarding the supernatant completely, 60 µL of separation buffer was added, followed by 40 µL of Anti-Biotin MicroBeads UltraPure (#130-105-637, Miltenyi Biotec, Bergisch Gladbach, Germany). The suspension was incubated for 15 minutes. The cells were then washed with 5 mL of separation buffer and resuspended in 500 µL of separation buffer, before proceeding to the magnetic separation step. The magnetic separation column, pre-filled with a MACS separator, was placed in a magnetic field. The column was rinsed with PBS before the cell suspension was added dropwise. Unlabeled cells were collected, and the column was washed 3 times, ensuring the column reservoir was empty each time. The column was then removed from the separator and placed in a collection tube. To elute the labeled cells, 1 mL of separation buffer was added to the column, and the plunger was applied firmly. The cell suspension was centrifuged and cultured in suspension flasks (T25, T75, or T175) pre-coated with Geltrex. Culture media consisted of DMEM F12 medium with 5 μg/mL insulin, 5 μg/mL transferrin, 5 μg/mL selenium, 35 ng/mL hydrocortisone, 10 ng/mL epidermal growth factor (EGF), 40 pg/mL triiodothyronine, and 2% fetal calf serum (FCS), supplemented with 1:100 Revitacell. After a maximum of 24 hours, Revitacell was removed and replaced with fresh medium. Cultures were maintained at 37 °C in a humidified incubator with 5% CO₂. Medium changes were performed every 2 or 3 days. Cells were passaged upon reaching ∼80% confluency. Cultures were maintained up to passage 3, with cells directly sorted from organoids considered to be passage 0.

### 5.3 Gentamicin exposure

The kidney organoids and LTL+ cells were exposed to gentamicin (gentamicin sulfate, #1405-41-0, Sigma-Aldrich, St. Louis, MO, USA) at a working concentration of 200mM in sterile miliQ. Treatment was applied to organoids on the days of medium refreshment and the following concentrations of gentamicin in appropriate medium were used: untreated, 0 mM, 0.25 mM, 0.5 mM, 1.0 mM, and 1.5 mM. Organoids were imaged under brightfield microscope every treatment day and harvested at d7+18. The LTL+ cells were exposed to gentamicin over 24 hours after reaching ∼80% confluency at passage 3 using the same concentrations. They were harvested after 24 hours of exposure for further processing.

### 5.4 Real Time Quantitative Polymerase Chain Reaction (RT-qPCR)

For each biological replicate four continuously treated organoids were collected at the end of the maturation protocol (d7+18) for qPCR analysis. First, the organoids were washed twice with PBS and then 300µL of lysis buffer (Buffer RLT provided with RNeasy Mini Kit) was added to lysate the cells. The cell lysate was stored at -20°C until RNA purification but no longer than 24 hours. As for the LTL+ cells, ∼145k cells at passage 3, cultured in a 12-well plate were collected after 24h of gentamicin exposure and the same protocol was used for the cell lysis as described for the organoids. RNA extraction was done using RNeasy Mini Kit (cat. Nos. 74104 and 74106) from Qiagen according to the manufacturer’s protocol. RNA concentration and quality was measured in Nanodrop. The cDNA was synthesized by incubating 600ng of RNA, 4µl of iScript RT Supermix, and nuclease-free water at a total volume of 20ul. Incubation was performed by priming for 5 minutes at 25 °C, performing reverse transcription for 20 minutes at 46 °C, and inactivating RT for 1 minute at 95 °C. The resulting cDNA was diluted ten times in nuclease-free water prior to qPCR. Lastly, RT-qPCR was performed using 2x SYBR Green Supermix and 10mM of each gene-specific forward and reverse primer. Reactions were carried out on the CFX96 Touch Real-Time PCR Detection System and CFX Opus 384 Real-Time PCR System using the following cycling conditions: initial denaturation for 3 minutes 95 °C; then 35 cycles of 5 seconds at 98 °C, 10 seconds at 60 °C and 20 seconds at 72 °C; 2 minutes at 72 °C and hold at 4 °C. The messenger RNA (mRNA) expression of each gene was normalized to the housekeeping gene HPRT1, and relative expression levels were calculated using the 2**^−ΔΔCT^** method.

### 5.5 LC-MS/MS

For each biological replicate, four organoids at d7+18 were pooled and washed twice with ice-cold PBS. Next the organoids were dissociated into single cells by incubation with pre-warmed trypsin (37°C) until complete disintegration was observed. Trypsinization was quenched by adding cold DMEM, and the cell suspension was centrifuged at 6000 rcf for 4 minutes at 4°C. After centrifugation, the supernatant was discarded, and the cell pallet was resuspended in 100µl of ice-cold methanol: MiliQ water (80:20, v/v). Samples were stored at -20°C for 24 hours. Then, the samples were vortexed until no cell debris remained visible, then centrifuged a 20 000 rcf for 4 minutes at 4°C. The supernatant was collected for LC-MS analysis, while the pellet - for BCA protein quantification. Around 145k of LTL+ cells at passage 3, cultured in a 12-well plate were lysed for further LC-MS analysis following the same protocol as explained above for the organoids. For LC-MS/MS analysis, compounds were separated using an Acquity UPLC® C18 column (30×2.1 mm, dp = 1.7 μm, Waters, Milford, USA) protected by an UPLC® BEH C18 VanGuard pre-column (5 ×2.1 mm, dp = 1.7 μm, Waters). The mobile phase consisted of (A) 0.1% formic acid with 10mM pentafluoropropionic acid in water and (B) 1% (v/v) dimethylsulfoxide in methanol, the total flow rate was 0.5 ml/min. Compounds were detected using AB Sciex QTRAP® 5500 (Ontario, Canada) triple quadrupole mass spectrometer equipped with a Turbo V™ TurboIonSpray® probe and an inlet valve. Key parameters for selected reaction monitoring detection included curtain gas (CUR) 20psi, collision gas (CAD) medium, ion spray voltage (IS) 3000V, temperature 750°C, ion source gas 1 (GS1) 70 psi, ion source gas 2 (IS2) 90psi, entrance potential (EP) 10V, collision cell exit potential (CXP) 13V. Stock solutions of gentamicin sulfate and kanamycin B (internal standard (IS)) were prepared in methanol-water-formic acid (50/50/0.1 (v/v)) at 1 mg/ml. Calibration standards (50–10,000 ng/ml) and quality control samples (160, 800, 8000 ng/ml) were prepared in appropriate matrices and stored at −30°C. For sample preparation, 10 µl of sample was mixed with 100 µl of internal standard solution (100 ng/ml kanamycin B in 0.1% formic acid water), vortexed, centrifuged, and 5 µl was injected for analysis.

### 5.6 Immunostainings

The kidney organoids cultured on Transwell inserts were collected at the end of the maturation protocol (d7+18) by excising the Transwell membranes with a sterile scalpel. Samples were washed twice with PBS and incubated in 4% paraformaldehyde (PFA) for 20 minutes at RT. After fixation, PFA was removed, and PBS was added to the samples for storage at 4C until further processing. For the LTL+, the cells cultured in 96-well plates were fixed using the same protocol as described for the organoids.

### 5.7 Organoid cryosectioning

Prior to embedding, fixed organoids were immersed in a 30% sucrose solution (w/v in PBS) and stored at 4°C for 24 hours before sectioning. Next, the sucrose solution was removed, and organoids were embedded in Tissue-Tek^®^ O.C.T. Compound (OCT, #4583, Sakura Finetek USA, Torrance, CA, USA). Embedded samples were sectioned using a cryostat at -18°C. Initial trimming was performed at 30mM thickness to reach the region containing the organoids. Once organoids were identified within the block, serial sections were collected at 5mM thickness onto microscope slides. Slides were dried overnight at RT and then stored at -80°C until further analysis. Cryostat sections were rinsed in phosphate-buffered saline (PBS) for 10 minutes and subsequently dried on a hot plate. Organoids were circled using a water-resistant marker. Blocking was performed for 2 hours at RT using 10% donkey serum in PBS, excluding Triton X-100 to preserve the integrity of the marker boundary. Cryosections were stained for the following biomarkers: kidney injury molecule-1 (Kim-1), collagen I, lotus tetragonolobus lectin (LTL), low-density lipoprotein receptor-related protein 2 (lrp2), and organic cation transporter 2 (OCT2). Primary antibodies were diluted in blocking buffer and incubated overnight at 4°C in a humidified chamber. Following primary antibody incubation, slides were washed three times with PBS. Secondary antibodies, also diluted in blocking buffer, were applied for 1 hour at RT followed by three additional PBS washes (3 minutes each). Nuclear staining was performed on the day of imaging using Hoechst 33342 (#62249, ThermoFisher, Waltham, MA, USA) at a dilution of 1:10,000 in PBS, incubated for 15 minutes. Slides were then washed three times for 3 minutes in PBS. A drop of Dako Fluorescence Mounting Medium (#S3023, Agilent Technologies, Santa Clara, CA, USA) was applied directly onto the organoids, and samples were sealed with coverslips. Imaging was performed at confocal microscope Leica TCS SP8 X.

### 5.8 Whole-organoid immunostaining

Organoids previously fixed in 4% paraformaldehyde (PFA) were blocked with 250 µL of 10% donkey serum in PBS containing Triton X-100 for 2-3 hours at RT. Following blocking, organoids were incubated with primary antibodies for 48 hours at 4 °C under gentle agitation in a cold room, each diluted 1:300 in blocking buffer, to visualize distinct nephron segments. Nephrin antibody was used to visualize glomeruli, LTL was used to visualize proximal tubules, E-cadherin - distal tubules, and GATA-3 - distal parts of the distal tubules and collecting ducts. Next, samples were washed three times with PBS for 10 minutes each. Secondary antibodies (look up section 2.1. Reagents and antibodies), diluted 1:400 in PBS containing 0.2% Triton X-100, were applied for 24 hours at 4 °C. After secondary antibody incubation, organoids were washed three times for 10 minutes each in PBS, transferred to Wilco dishes, and mounted using Dako Fluorescence Mounting Medium. Imaging was performed using a Leica TCS SP8 X confocal microscope.

### 5.9 LTL+ immunostaining

The fixed LTL+ were first blocked with donkey serum-based blocking buffer for 1.5 hours at RT on a gentle shaker. Primary antibodies diluted in blocking buffer were added and incubated for 1.5 hours under gentle shaking. Following three PBS washes (≥5 minutes each), the secondary antibodies diluted in PBS with 0.2% Triton X-100 were added for 1.5 hours under gentle shaking, protected from light. After three additional PBS washes, plates were stored at 4 °C in PBS, covered with foil. On the day of imaging, Hoechst 33342 (1:10,000 in PBS) was added to each well and incubated for 15 minutes at RT. Wells were washed three times with PBS and replenished with PBS prior to imaging. Antigens and antibodies are depicted in the table below (Table 1).

**Table 1.** Antibodies with their dilutions used for fixed immunostainings of LTL+ cells.

| **Antigen** | **Primary antibody** | **Dilution** | **Secondary antibody/ probe** | **Dilution** |
|---|---|---|---|---|
| KIM-1 | TIM-1 Polyclonal antibody, Invitrogen, #PA5-20244 | 1:25 | Donkey anti-rabbit 647, Invitrogen, #A31573 | 1:400 |
| BODIPY | - | - | BODIPY 493/ 503 10mM | 1:1000 |
| LTL | Biotinylated Lotus Tetragonolobus Lectin, Vector Laboratories, Brunschwig Chemie, #B-1325 | 1:100 | Streptavidin Alexa Fluor 405 conjugate, Thermo Fisher Scientific, #S32351 | 1:400 |
| Megalin | Mouse anti-LRP2/ megalin Sigma Aldrich | 1:100 | Donkey anti-rabbit 647, Invitrogen, #A31573 | 1:400 |
| Phalloidin | - | - | Phalloidin 647 | 1:1000 |

### 5.10 Azan Trichrome staining

Azan Trichrome staining was performed on fixed cryosections of kidney organoids at d7+18. Azan Trichrome was performed following the methods provided in the Azan Trichrome kit (#04-001802, Bio-Optica, Milan, Italy). In this method two acid dyes were used, azocarmine (pink) which stains non-collagenous tissue, and aniline blue (blue) which stains collagenous tissue.

### 5.11 Live immunostainings in LTL+ cells

The LTL+ cells (passage 3) were seeded into 96-well Yokogawa plates pre-coated with 2% Geltrex in DMEM/F12 for 1 hour. Cells were plated 24 hours prior to staining. For nuclear labeling, cells were incubated with Hoechst (1:10,000) for 15 minutes at 37 °C. Subsequently, gentamicin and fluorescent probes were added at the following concentrations: LysoTracker Green DND-26 50 nM (#L7526, Invitrogen, Waltham, MA, USA), BODIPY 581/591 lipid peroxidation 10 mM (#D3861, ThermoFisher, Waltham, MA, USA). Imaging was initiated 30 minutes after probe addition to allow for the probe’s incubation described by the manufacturer. Plates were imaged on the Yokogawa high-content imaging system for 24 hours, with images acquired every hour. Throughout the experiment, exposure time, laser power, and z-stack parameters were kept constant.

### 5.12 Image analysis and statistical analysis

Image analysis was done in Fiji (ImageJ 2.16.0, National Institutes of Health, Bethesda, MD, USA). Thresholds were determined based on auto threshold function and kept constant across all treatment conditions for each staining. Images of the whole organoids were rendered and subsequently analyzed in IMARIS 10.2 (Oxford Instruments/Andor, Zurich, Switzerland). An identical threshold diameter of 40mM was applied across all treatment groups to define individual tubules. Results were depicted as the mean plus-or-minus (±) standard error of mean (SEM). Statistical analysis was conducted in GraphPad Prism 10 (GraphPad software, La Jolla, CA, USA). Intracellular uptake of gentamicin was analyzed using a two-way mixed effects model. Organoid growth was analyzed using Friedman test with Dunn’s multiple comparisons test (vs. untreated). The rest of data gathered was analyzed using a One-Way ANOVA Kruskal-Wallis test followed by Dunn’s multiple comparisons. A p-value of < 0.05 was considered as statistically significant. Every experiment was performed in at least three biological replicates, including at least 2 technical replicates, unless specified otherwise. Each biological replicate represents an organoid derived from a different differentiation. Each technical replicate represents either a different organoid within the same differentiation batch or a different slice from the same organoid. For LTL+ cells, each biological replicate represents LTL+ cells sorted from organoids of separate differentiation runs. Technical replicates were obtained from different wells of the same differentiation experiment.

## 6. AUTHOR CONTRIBUTIONS

E.S.G. and A.M.G. designed the *in vitro* study. E.S.G., W.P., L.B. and R.W.S. performed the experiments. E.S.G., W.P., L.B. and R.W.S. analyzed the data. E.S.G. wrote the manuscript. A.M.G. and R.M. reviewed and edited the manuscript. All authors read and contributed to the final version of the manuscript.

### Ethics approval and consent to participate

Human erythroblasts, after giving informed consent, were used to generate iPSCs. This study was conducted in accordance with the Helsinki Declaration as revised in 2013. Permission for the creation and use of iPSCs in this study was obtained from the local ethical commission for human-related research of the Radboud University Medical Center, Nijmegen (2015-1543).

### Data availability statement

The raw data supporting the conclusions of this article will be made available by the authors upon reasonable request.

## Acknowledgments

This work was supported by the Virtual Human Platform for Safety Assessment project, which is funded by the Netherlands Research Council (NWO) ‘Netherlands Research Agenda: Research on Routes by Consortia’ (NWA ORC 1292.19.272).

## REFERENCES

Allegaert K (2009) Aminoglycoside pharmacokinetics in neonates: the need to search for covariates of variability in addition to case descriptions. Acta Paediatr 98(10):1546; author reply 1546-7 doi:10.1111/j.1651-2227.2009.01420.x

Balakumar P, Rohilla A, Thangathirupathi A (2010) Gentamicin-induced nephrotoxicity: Do we have a promising therapeutic approach to blunt it? Pharmacol Res 62(3):179–86 doi:10.1016/j.phrs.2010.04.004

Beenken A, Cerutti G, Brasch J, et al. (2023) Structures of LRP2 reveal a molecular machine for endocytosis. Cell 186(4):821–836.e13 doi:10.1016/j.cell.2023.01.016

Berrout J, Jin M, Mamenko M, Zaika O, Pochynyuk O, O’Neil RG (2012) Function of transient receptor potential cation channel subfamily V member 4 (TRPV4) as a mechanical transducer in flow-sensitive segments of renal collecting duct system. J Biol Chem 287(12):8782–91 doi:10.1074/jbc.M111.308411

Bulman ZP, Cirz R, Hildebrandt D, et al. (2020) Unraveling the Gentamicin Drug Product Complexity Reveals Variation in Microbiological Activities and Nephrotoxicity. Antimicrob Agents Chemother 64(9) doi:10.1128/AAC.00533-20

Chattopadhyay B (2002) Newborns and gentamicin--how much and how often? J Antimicrob Chemother 49(1):13–6 doi:10.1093/jac/49.1.13

Choi JS, Kim CK, Lee BJ (1999) Administration-time differences in the pharmacokinetics of gentamicin intravenously delivered to human beings. Chronobiol Int 16(6):821–9 doi:10.3109/07420529909016948

Dagil R, O’Shea C, Nykjær A, Bonvin AM, Kragelund BB (2013) Gentamicin binds to the megalin receptor as a competitive inhibitor using the common ligand binding motif of complement type repeats: insight from the nmr structure of the 10th complement type repeat domain alone and in complex with gentamicin. J Biol Chem 288(6):4424–35 doi:10.1074/jbc.M112.434159

Denamur S, Tyteca D, Marchand-Brynaert J, et al. (2011) Role of oxidative stress in lysosomal membrane permeabilization and apoptosis induced by gentamicin, an aminoglycoside antibiotic. Free Radic Biol Med 51(9):1656–65 doi:10.1016/j.freeradbiomed.2011.07.015

Fjalstad JW, Laukli E, van den Anker JN, Klingenberg C (2013) High-dose gentamicin in newborn infants: is it safe? Eur J Pediatr doi:10.1007/s00431-013-2194-1

Gilbert T, Lelievre-Pegorier M, Merlet-Benichou C (1990) Immediate and long-term renal effects of fetal exposure to gentamicin. Pediatr Nephrol 4(4):445–50 doi:10.1007/BF00862534

Guo JK, Menke AL, Gubler MC, et al. (2002) WT1 is a key regulator of podocyte function: reduced expression levels cause crescentic glomerulonephritis and mesangial sclerosis. Hum Mol Genet 11(6):651–9 doi:10.1093/hmg/11.6.651

Hodiamont CJ, van den Broek AK, de Vroom SL, Prins JM, Mathôt RAA, van Hest RM (2022) Clinical Pharmacokinetics of Gentamicin in Various Patient Populations and Consequences for Optimal Dosing for Gram-Negative Infections: An Updated Review. Clin Pharmacokinet 61(8):1075–1094 doi:10.1007/s40262-022-01143-0

Hostetler KY, Hall LB (1982) Inhibition of kidney lysosomal phospholipases A and C by aminoglycoside antibiotics: possible mechanism of aminoglycoside toxicity. Proc Natl Acad Sci U S A 79(5):1663–7 doi:10.1073/pnas.79.5.1663

Keller S, Luciani A, Devuyst O (2024) The structure of megalin: shedding new light on receptor-mediated endocytosis. Kidney Int 105(1):11–14 doi:10.1016/j.kint.2023.05.026

Klementa V, Petejova N, Horak P, Kurasova E, Zadrazil J (2025) Acute kidney injury due to gentamicin nephrotoxicity and specific miRNAs as biomarkers. Biomed Pap Med Fac Univ Palacky Olomouc Czech Repub 169(1):1–8 doi:10.5507/bp.2024.031

Kobayashi A, Valerius MT, Mugford JW, et al. (2008) Six2 defines and regulates a multipotent self-renewing nephron progenitor population throughout mammalian kidney development. Cell Stem Cell 3(2):169–81 doi:10.1016/j.stem.2008.05.020

Lei L, Li LP, Zeng Z, et al. (2018) Value of urinary KIM-1 and NGAL combined with serum Cys C for predicting acute kidney injury secondary to decompensated cirrhosis. Sci Rep 8(1):7962 doi:10.1038/s41598-018-26226-6

Lingvall M, Reith D, Broadbent R (2005) The effect of sepsis upon gentamicin pharmacokinetics in neonates. Br J Clin Pharmacol 59(1):54–61 doi:10.1111/j.1365-2125.2005.02260.x

Lulic-Botica M, Sheer T, Edwards D, Thomas RL, Natarajan G (2014) Impact of small-for-gestational age (SGA) status on gentamicin pharmacokinetics in neonates. J Clin Pharmacol 54(1):39–45 doi:10.1002/jcph.190

Mingeot-Leclercq MP, Tulkens PM (1999) Aminoglycosides: nephrotoxicity. Antimicrob Agents Chemother 43(5):1003–12 doi:10.1128/AAC.43.5.1003

Nagai J, Takano M (2014) Entry of aminoglycosides into renal tubular epithelial cells via endocytosis-dependent and endocytosis-independent pathways. Biochem Pharmacol 90(4):331–7 doi:10.1016/j.bcp.2014.05.018

Nakanoh H, Tsuji K, Fukushima K, et al. (2025) Kidney Organoids: Current Advances and Applications. Life (Basel) 15(11) doi:10.3390/life15111680

Nguyen J, Madonia V, Bland CM, et al. (2025) A review of antibiotic safety in pregnancy-2025 update. Pharmacotherapy 45(4):227–237 doi:10.1002/phar.70010

Phillips I, Eykyn S, King BA, Jenkins C, Warren CA, Shannon KP (1977) The in vitro antibacterial activity of nine aminoglycosides and spectinomycin on clinical isolates of common Gram-negative bacteria. J Antimicrob Chemother 3(5):403–10 doi:10.1093/jac/3.5.403

Quiros Y, Vicente-Vicente L, Morales AI, López-Novoa JM, López-Hernández FJ (2011) An integrative overview on the mechanisms underlying the renal tubular cytotoxicity of gentamicin. Toxicol Sci 119(2):245–56 doi:10.1093/toxsci/kfq267

Randjelovic P, Veljkovic S, Stojiljkovic N, Sokolovic D, Ilic I (2017) Gentamicin nephrotoxicity in animals: Current knowledge and future perspectives. EXCLI J 16:388–399 doi:10.17179/excli2017-165

Rodríguez Salgueiro S, González Núñez L (2016) Animal models mimicking aminoglycoside-induced renal damage. J Nephropharmacol 5(1):1–3

Serane TV, Zengeya S, Penford G, Cooke J, Khanna G, McGregor-Colman E (2009) Once daily dose gentamicin in neonates - is our dosing correct? Acta Paediatr 98(7):1100–5 doi:10.1111/j.1651-2227.2009.01297.x

Wang G, Wu H, Zhai X, et al. (2024) Kidney Organoid Modeling of WT1 Mutations Reveals Key Regulatory Paths Underlying Podocyte Development. Adv Sci (Weinh) 11(29):e2308556 doi:10.1002/advs.202308556

Wargo KA, Edwards JD (2014) Aminoglycoside-induced nephrotoxicity. J Pharm Pract 27(6):573–7 doi:10.1177/0897190014546836

Watanabe M (1978) Drug-induced lysosomal changes and nephrotoxicity in rats. Acta Pathol Jpn 28(6):867–89 doi:10.1111/j.1440-1827.1978.tb01277.x

Wells VD, Wong ES, Murray BE, Coudron PE, Williams DS, Markowitz SM (1992) Infections due to beta-lactamase-producing, high-level gentamicin-resistant Enterococcus faecalis. Ann Intern Med 116(4):285–92 doi:10.7326/0003-4819-116-4-285

Yoshimura Y, Muto Y, Omachi K, Miner JH, Humphreys BD (2023) Elucidating the Proximal Tubule HNF4A Gene Regulatory Network in Human Kidney Organoids. J Am Soc Nephrol 34(10):1672–1686 doi:10.1681/ASN.0000000000000197

